# Physiological Gaussian Process Priors for the Hemodynamics in fMRI Analysis

**DOI:** 10.1101/179838

**Authors:** Josef Wilzén, Anders Eklund, Mattias Villani

**Author notes:** Corresponding author Email address (Josef Wilzén*).

## Abstract

Inference from fMRI data faces the challenge that the hemodynamic system, that relates the underlying neural activity to the observed BOLD fMRI signal, is not known. We propose a new Bayesian model for task fMRI data with the following features: (i) joint estimation of brain activity and the underlying hemodynamics, (ii) the hemodynamics is modeled nonparametrically with a Gaussian process (GP) prior guided by physiological information and (iii) the predicted BOLD is not necessarily generated by a linear time-invariant (LTI) system. We place a GP prior directly on the predicted BOLD time series, rather than on the hemodynamic response function as in previous literature. This allows us to incorporate physiological information via the GP prior mean in a flexible way. The prior mean function may be generated from a standard LTI system, based on a canonical hemodynamic response function, or a more elaborate physiological model such as the Balloon model. This gives us the nonparametric flexibility of the GP, but allows the posterior to fall back on the physiologically based prior when the data are weak. Results on simulated data show that even with an erroneous prior for the GP, the proposed model is still able to discriminate between active and non-active voxels in a satisfactory way. The proposed model is also applied to real fMRI data, where our Gaussian process model in several cases finds brain activity where previously proposed LTI models, parametric and nonparametric, does not.

## 1. Introduction

### 1.1. Background

Task based fMRI data are typically analyzed using voxel-wise general linear models (GLM), to detect voxels or regions where the blood oxygenation level dependent (BOLD) contrast is correlated with the experimental stimuli paradigm (Friston et al., 1994; Lindquist et al., 2008). BOLD is an indirect measure of neural activation which depends on the hemodynamic response (HR). Understanding the HR is therefore critical in order to correctly infer the brain activity (Handwerker et al., 2004; Lindquist and Wager, 2007; Lindquist et al., 2009). The neurovascular coupling between the neural response triggered by a stimulus and the observed BOLD response in fMRI is not fully understood (Logothetis, 2002, 2003), and the HR has been shown to vary across voxels, brain regions and subjects (Handwerker et al., 2004, 2012). It is common practice in fMRI to model the HR as a linear time invariant system (LTI) (Boynton et al., 2012). Standard GLMs make very strong assumptions about the HR, and since it is unlikely that these models are correct for all voxels and subjects, the inference for the brain activity parameters will be biased (Lindquist and Wager, 2007).

### 1.2. Joint Detection Estimation framework and Gaussian Processes

The so called joint detection estimation (JDE) framework for the GLM estimates the brain activity jointly with the HR. The JDE approach uses a zero mean Gaussian process prior on filter coefficients, which represent the HR in a LTI context. The filter is often called the hemodynamic response function (HRF) (Goutte et al., 2000; Ciuciu et al., 2003; Marrelec et al., 2003a; Casanova et al., 2008). The activation strength for each voxel is based on a summary statistics on the whole filter. A problem with such voxel-wise approaches is that the filter is unidentified if the specific voxel is inactive. There is also a high risk of overfitting, since a separate HR is estimated in each voxel.

Another approach is to use a bilinear model where both the design matrix and the regression coefficients are estimated jointly. Many models based on the JDE framework use parcellation, see for example Makni et al. (2008); Vincent et al. (2010). Some parameters are constant within each parcel, while other parameters are voxel specific. Parcellation can be done a priori and considered constant (Makni et al., 2008; Vincent et al., 2010), or estimated as a part of the model (Chaari et al., 2012, 2016; Albughdadi et al., 2016). The idea is to restrict the HR to be the same across all voxels in a parcel, but allowing the activation parameters to vary across the voxels within a parcel. In the joint parcellation-detection-estimation (JPDE) framework, the parcellation itself is also learned from the data (Chaari et al., 2012, 2016; Albughdadi et al., 2016). These models use a random effect approach, where the HRF for a given voxel is a random draw from a distribution with a parcel-specific mean HRF. The proposed JDE and JPDE models have been analyzed by variants of Gibbs sampling (Makni et al., 2005, 2006b,a; Ciuciu et al., 2007; Vincent et al., 2007; Makni et al., 2008; Ciuciu et al., 2009; Vincent et al., 2010) and the approximate, but quicker, variational Bayes (VB) method (Chaari et al., 2011, 2012, 2013, 2016; Albughdadi et al., 2016).

Different assumptions have been made regarding the Gaussian process prior for the HRF in the JDE literature. In order to assume a causal filter, the endpoints of the filter are often constrained to be zero (Goutte et al., 2000; Marrelec et al., 2003b; Ciuciu et al., 2003; Makni et al., 2005; Ciuciu et al., 2007; Vincent et al., 2007; Makni et al., 2008; Ciuciu et al., 2009; Vincent et al., 2010; Chaari et al., 2011, 2012, 2013, 2016; Albughdadi et al., 2016). Goutte et al. (2000) and Quirós et al. (2010) use a squared exponential kernel; others use the second order difference matrix as precision matrix (Marrelec et al., 2003b; Ciuciu et al., 2003; Makni et al., 2005, 2006b,a; Ciuciu et al., 2007; Vincent et al., 2007; Makni et al., 2008; Ciuciu et al., 2009; Vincent et al., 2010) and variants thereof (Chaari et al., 2011, 2012, 2013, 2016; Albughdadi et al., 2016). For the approaches that use a fixed and a priori known parcellation, it is assumed that there is one HRF per parcel, regardless of the number of stimuli in the experiment. The idea is that a functionally similar region has the same hemodynamic behavior.

### 1.3. Non-linearity of Predicted BOLD

There is, however, evidence that contradicts the LTI system hypothesis for the HR, see for example Huettel et al. (2004) for a discussion. This has motivated the development of more physiologically realistic models that do not assume an LTI system, and model the predicted BOLD directly (Buxton et al., 1998; Friston et al., 1998, 2000; Buxton et al., 2004; Deneux and Faugeras, 2006; Stephan et al., 2007; Lundengàrd et al., 2016). Estimation of such nonlinear models are typically more computationally expensive. Non-linear extensions of JDE models that focus on the non-linear habituation effect of repeated stimuli (Ciuciu et al., 2009) are more efficient, but accounts only for a limited class of non-linearities.

### 1.4. Our approach

In this work, we propose a new model that places a Gaussian Process (GP) prior (Rasmussen and Williams, 2006) directly on the predicted BOLD time series. This is in contrast to earlier work which use a Gaussian process prior on the HRF, and then convolve the posterior HRF with the paradigm. Our approach is therefore not restricted to LTI systems, which means that non-stationary and non-linear properties of the BOLD response can be handled, if supported by the data. Non-stationarity of the BOLD response can for example arise from refractory and adaptation effects (Huettel et al., 2004), or from a participant:s failure to perform a task in the MR scanner. Our approach can also implicitly account for the so called stimulus-as-fixed-effect fallacy (Westfall et al., 2016).

A GP prior on the predicted BOLD makes the model very flexible, which can lead to overfitting. Our model therefore incorporates several features to avoid overfitting. First, we use a parcellation approach similar to Makni et al. (2008); Vincent et al. (2010), where the predicted BOLD is restricted to be the same for all voxels in a given parcel, but the activation and other parameters (for example modeling time trends) are voxel-specific. The effect is that the predicted BOLD in a parcel is accurately estimated from data in many voxels. Second, in contrast to the JDE literature, the mean of our GP prior is non-zero and equal to the predicted BOLD from a physiologically motivated model of the hemodynamics, for example the Balloon model proposed by Buxton et al. (1998, 2004). This allows the GP posterior to fall back on the baseline physiological model whenever the data are weak or support the baseline model, while still being able to override the prior mean when the data are incompatible with the baseline model. Third, using a well founded prior mean makes it possible to use relatively tight GP kernels.

The rest of the paper is organized as follow: Section 2 describes the model and the prior on its parameters. The inference procedure is presented in Section 3. Results from simulations and real data are given in Section 4 and 5, respectively. The paper ends with a discussion in Section 6 and conclusions in Section 7.

## 2. Model, Priors and Posterior Computations

### 2.1. Notation

Vectors and matrices are denoted with bold lower and upper case letters, respectively. Vectors are assumed to be column vectors. The symbol *^T^* denotes transposeп*_a_* denotes the identity matrix of size *a × a*, *vec*(.) is the vectorization operator ⊗, is the Kronecker product and *diag*(*x*) means a diagonal matrix with vector *x* as the main diagonal. *N* (*μ*, **Σ**) and *M N* (*μ*, **Σ**, **Ω**) denote multivariate normal and matrix normal (see Appendix A) distributions, respectively. *InvGamma*(*a, b*) denotes the inverse gamma distribution. The following different indices are used:

- *j*: voxels, *j* ∈ {1*, …, J*}, within a parcel
- *m*: stimulus type, *m* ∈ {1*, …, M*}
- *p*: number of nuisance variables, *p* ∈ {1*, …, P*}
- *k*: number of parameters in the *AR*(*k*) process, *k* ∈ {1*, …, K*}.
- *t*: time, 
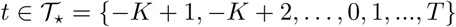
.

### 2.2. A physiological Gaussian process prior for predicted BOLD

The fMRI signal will be modeled in the following way: hemodynamic responses are the same for all voxels in a parcel, while task related activations and parameters for the noise process are allowed to vary between voxels in a parcel. The time series contain temporal autocorrelation, which in our case is modeled using an autoregressive (AR) process of order *K*. We make the usual simplifying assumption in time series analysis that the first *K* values of the process are known. Let 
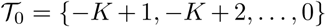
 be an initial set of time points where the data values are assumed to be known. Further, define 
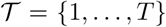
 to be the subsequent time points and 
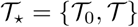
 to be the set of all *T*_*_ = *T* + *K* time points.

The predicted BOLD is modeled with a Gaussian process prior (Rasmussen and Williams, 2006) according to

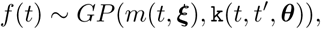

where *m*(*t*, *ξ*) is the mean function and *k*(*t*, *t*′, *θ*) is the kernel (covariance function) of the process. A sampled value of the GP is denoted *f_t_* and

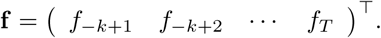

This prior gives a general framework for modeling the hemodynamic response with a variety of physiological models or constraints. The mean function, defined by the parameters *ξ*, can come from some arbitrary model that can generate the predicted BOLD. These models can be linear (e.g. the HRF used in the SPM software) or nonlinear (e.g. the Balloon model). The kernel controls both the degree of smoothness of **f** and the deviation from the mean function. If several stimuli are considered, the total predicted BOLD is considered to be a linear combination of different GPs, which will be denoted **f***_m_*, and the(*T* + *K*) × *m* matrix **F** = (**f_1_ … f_*M*_**) gather all the sampled realizations of the different GPs. The parameters *ξ*_*m*_ and *θ_m_* denote the stimulus specific GP hyperparameters. We use a Matèrn kernel with 5/2 degrees of freedom

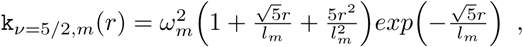

where *r* is the Euclidean distance between two covariate observations. Let 
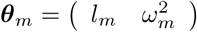
 and *θ* = (*θ*_1_, …, *θ_M_*). The covariance matrix for all data points from k(*t, t′*, *θ_m_*)) is denoted 
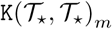
.

The voxel and stimulus specific activations will be represented by the parameter *β_m,j_* which is contained in the matrix **B**, of size *M × J*, where *J* is the number of voxels. One approach is to let the voxel-wise hemodynamics be modeled as **FB**. The drawback with this approach is that the parameters enter the likelihood as a product, and are thereby not individually identified since **FB = F***S*^−1^*S***B**, for any invertible matrix *S* of size *M* × *M*. To overcome this problem, we propose an identifying nonlinear transformation for the matrix **F**. The transformation must fixate the scale of **F**, and prevent sign flipping as well as linear combinations and permutations. Similar to Pedregosa et al. (2015), we propose the transformation:

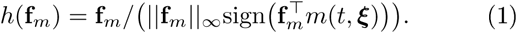

This transformation is still sensitive to pure permutations, so in order to identify the column order we introduce a permutation function Ψ(·). Let **F**_0_ be prior mean for **F** and **F***_post_* be a sample from the posterior for **F**. The function Ψ(**F***_post_*) permutes the column in **F***_post_* in such a way that

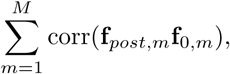

is maximized with respect to the column order of **F***_post_*. The final transformation is then given by

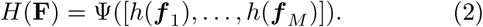

*H*(**F**) has its support on a relative scale and is bounded between -1 and 1.

### 2.3. Multivariate GLM for joint detection and estimation

We use a multivariate regression model for all observed BOLD signals in one parcel, i.e.

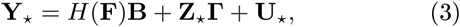

where **Y***_*_* = (**y**_1_ **y***_J_*) is a *T_*_* × *J* matrix with the observed BOLD signal, where columns represent different voxels and rows represent different time points. The matrix **Z***_*_*, which is of size *T_*_* × *P*, contains nuisance covariates, such as time trends and head motion covariates, and the corresponding regression coefficients are stored in the *P* × *J* matrix **Γ**. **U***_*_* = (**u**^(1)^ • • • **u**^(*J*)^) is the error matrix, of size *T_*_ × J*, which contains the corresponding errors for all elements in **Y**. The matrix **F** and its transformation *H*(**F**) are both of size *T_*_* × *M*. The idea is to let *H*(**F**) model the dynamics in the predicted BOLD, while **B** models the overall response magnitude in each voxel. The separation of the hemodynamics from the activations gives a straight forward measure of voxel activation, that can be used to construct posterior probability maps (PPMs) or *t*maps. The hyperparameters for the kernel are for simplicity assumed to be fixed and known, but can in principle be learned in separate updating steps; see the discussion in Section 5.5. The conditional dependency of *ξ* and *θ* are omitted in the rest of the paper for notational clarity.

Equation (3) can be partitioned with respect to the time observation that corresponds to the pre-sample observations in 
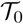
 and the estimation sample in 
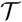
 respectively, according to

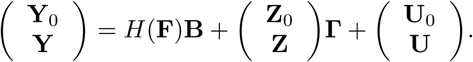

The upper parts of **Y***_*_* and **Z***_*_* have *K* rows, and the lower parts have *T* rows. The upper parts will be used as lags in different pre-whitening steps and the lower parts will be used for the inference.

We assume that the noise in each voxel follows an *AR*(*K*) process, i.e. for the *j*th voxel

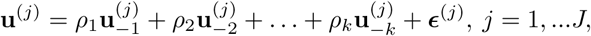

where the negative indices denote time lags. We assume that the AR parameters are the same for all voxels in a parcel, but different across parcels. The error terms *E*^(*j*)^ are assumed to be independent across voxels and 
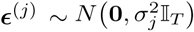
. Spatial noise dependencies can also be incorporated by replacing П*_T_* with a matrix **H** that models spatial dependences among the elements. The distribution of **U** can be expressed as

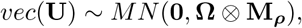

where **Ω** = *diag σ*^2^ and 
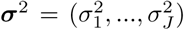
. The matrix **M**_*ρ*_ can be obtained by solving a system of Yule-Walker equations or using the methods of van der Leeuw (1994), but it is not needed explicitly for sampling from the posterior of the model in (3).

The user of our model must decide the parcel sizes. Larger parcels use more data for the estimation of the predicted BOLD, which will be more robust to noise and inactivity, but a drawback is a lower flexibility. Smaller parcels will provide a higher flexibility, at the risk of overfitting.

### 2.4. Likelihood function and priors

The likelihood function for the model in (3) is of the form

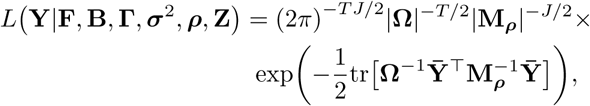

where **Y¯** = **Y** – *H*(**F**)**B** – **ZΓ**.

The model (3) has the following parameters and priors:

1. **F** has independent Gaussian process priors on each column, i.e.

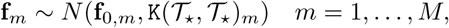

where **f**_0,*m*_ is the mean function *m*(*t*) evaluated at the time points 
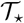
.
2. The elements of *σ*^2^ are assumed to be independent apriori and are modeled as

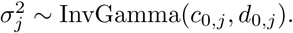
3. The prior for *B* is modeled conditional on **Ω** = diag(*σ*^2^) as a matrix normal distribution

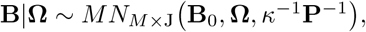

where **P** is a *M* × *M* positive definite precision matrix over stimuli, **B**_0_ is the *M × J* prior mean matrix and *κ* is a scalar.
4. **Γ** is assigned a matrix normal prior conditional on **Ω**

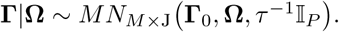
5. Following Eklund et al. (2016), the prior on the AR process parameters is centered over a stationary AR(1) process :

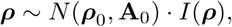

where *ρ*_0_ = (*r,* 0*, …,* 0), 
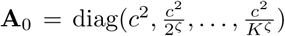
 and *I*(*ρ*) is an indicator function for the stationary region

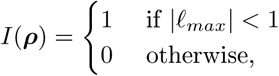

where *l_max_* is the largest absolute (modulus) eigenvalue of the companion matrix

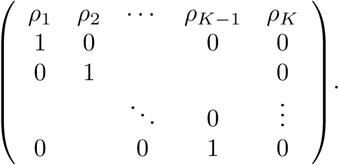 Note that the prior is centered over the noise process *u_t_* = *ρ*_1_.*u*_*t*−1_+*ε*_*t*_, but assigns probability mass also to higher order AR processes in such a way that longer lags are shrunk more heavily toward zero.

### 2.5. Posterior computations

The joint posterior of all model parameters in (3) is given by

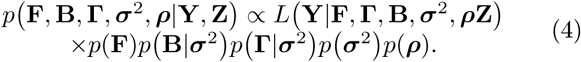

Equation (4) is intractable and cannot be sampled using standard distributions, and we therefore resort to Gibbs sampling. The posterior is sampled in four steps, which are described in Algorithm 1 and detailed formulas are given in the Appendix B.

### 2.6. Implementation

The proposed estimation of the model in (3) is implemented in the R programming language. Since each parcel is independent, the computation can be parallelized across parcels, which is done with the foreach package (Analytics and Weston, 2015). Functions from the R package neuRosim (Welvaert et al., 2011) are used to simulate data for the simulation study.

#### Algorithm 1 Schematic description of the Gibbs sampler for the posterior in Equation (4)

1. *ρ* is sampled from a multivariate Gaussian distribution in Equation (6). *ρ* is then used to pre-whiten the data.
2. *σ*^2^ is sampled from an Inverse-gamma distribution in Equation (9).
3. *vec*(*B*) and *vec*(**Γ**) are sampled from a multivariate normal distribution in Equation (10).
4. **F** is sampled with Elliptical slice sampling using the likelihood in (12).

## 3. Simulations

A simulation study was performed to investigate the proposed model:s ability to detect activity, and to estimate the underlying hemodynamics. Two models were used. In the first model, the predicted BOLD is fixated to the prior mean. For the second model, the predicted BOLD is instead estimated from the data using our physiological GP prior. Data were generated from the first model. A total of 32 simulated datasets were generated, for each combination of parameters in the data generating process. In each simulation a single parcel with 100 voxels was used, and 20 of the voxels were active. The following settings were used for each simulation

- A single stimulus was modeled as a block paradigm. The number of time points was set to 150 and the sampling rate (TR) was set to 1 second.
- An AR(3) process was used for the noise process, with autoregressive parameters: *ρ^T^* = (0.4, 0.1, 0.05).
- The contrast to noise ratio (CNR), *b_m,j_/σ*, where *b_m,j_ =* 1 for active voxels and zero otherwise, was set to 5 and 7.
- Constant, linear, quadratic and cubic trends were added to each time series. The coefficients were generated randomly for each voxel and simulation, to reflect realistic trends in fMRI data.

The same priors were used for all simulations, except for **Γ**, and were specified to:

1. **F**: The prior mean for **F** was created in such a way that is was either correct, with correlation 1 with the true process, or erroneous with a correlation of 0.615 with the true process. We use two different lengthscales: *l* = 2 or *l* = 4, and 
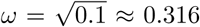
. The prior mean function was specified to the common double gamma HRF convolved with the paradigm. For the fix model **F** is not estimated, but instead fixated to the value of the prior mean.
2. *σ*^2^: *c*_0_ = 0 and *d*_0_ = 0, giving a non-informative prior.
3. **B**: precision scale factor *κ* = 10^−10^, precision matrix **P** is diagonal, giving an essentially flat noninformative prior.
4. **Γ**: *τ* = 0, except for the parameter representing the constant, which was given an empirical prior based on voxel mean and four times the voxel variance for each voxel. The nuisance variables are scaled to have zero mean and unit variance.
5. *ρ*: *ρ*_0_ = (0, 0, 0), 
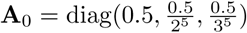

The posterior was sampled 4000 times and 1000 samples were discarded as burn in. The remaining samples were thinned out by a factor of 3, leaving 1000 posterior draws for inference.

There are several ways that voxels can be declared active in an Bayesian model. One way is to construct posterior probabilities of the type *p*(*b_m,j_* > *c* | **y**) > *a*, where *c* is a user defined effect size and *a* is a probability threshold. These probabilities can be used to construct PPMs over the brain. However, there is no consensus regarding how to threshold PPMs, and PPMs can be numerically unstable and imprecisely estimated. Instead, we use Bayesian “ *t*-ratios', defined as

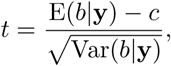

which can be easily computed from the posterior samples in each voxel. These ratios have a higher resolution for a fixed amount of posterior samples compared to PPMs. In the simulations *c* = 0 and the test *t > a* was used, where *a* is a quantile from a *t*-distribution. Note that the activation is voxel independent given **F** and *ρ*, due to the simulation design. These frequentist calculations are of course not directly transferable to a Bayesian setting, but has the advantage of giving familiar thresholds for fMRI. A range of values for *a* were tested and ROC curves were calculated. Figure 1 and 2 show the results for the lengthscales *l* = 4 and *l* = 2, respectively. The results show that our model with a GP prior on predicted BOLD model performs better for a wide range of thresholds, and thus has a much better ability to discriminate between active and non-active voxels. As expected, for very low thresholds the model with estimated predicted BOLD starts to classify non-active voxels as active.

Figure 3 and 4 show the posterior of the predicted BOLD for selected simulations.

## 4. Real data

### 4.1. Data

To test our proposed approach on real data, we used open fMRI data from brain tumor patients (Pernet et al., 2016; Gorgolewski et al., 2013), as the hemodynamic response function may be different close to a tumor. A total of 22 patients (9 females) with different types of brain tumors were scanned using both structural (T1, T2, DWI) and functional (BOLD T2*) MRI sequences. For the functional scans several tasks were performed: motor, verb generation and word repetition (resting state data are also available). Data were acquired on a General Electric 1.5 Tesla scanner with an 8 channel phased-array head coil. The fMRI data were acquired using a standard EPI sequence with a repetition time of 5.0 seconds (due to sparse sampling for auditory tasks) or 2.5 seconds, and an echo time of 50 milliseconds. Each voxel has a size of 4 × 4 × 4 mm^3^, resulting in volumes with 64 × 64 × 30 voxels.

**Figure 1:**
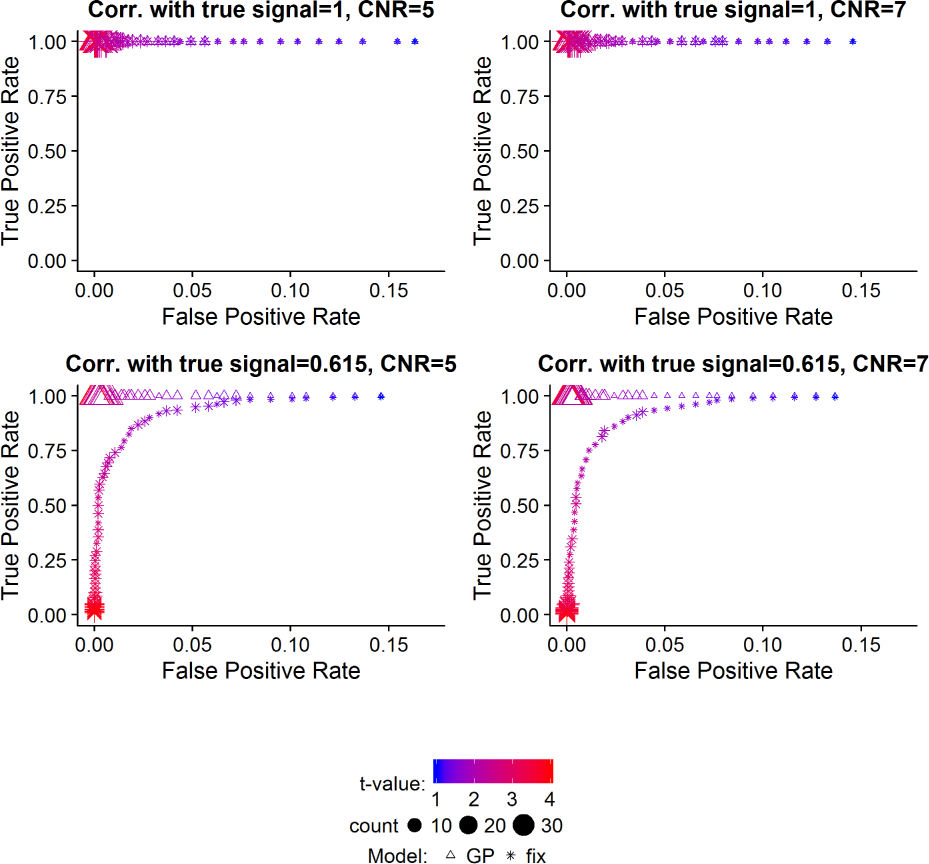
ROC curves for the simulation study. Lengthscale hyperparameter for GP: *l* = 4. Each value of is the average of over 32 simulations. Thresholds for *t*-values: 60 equidistant values between 1 and 4.

**Figure 2:**
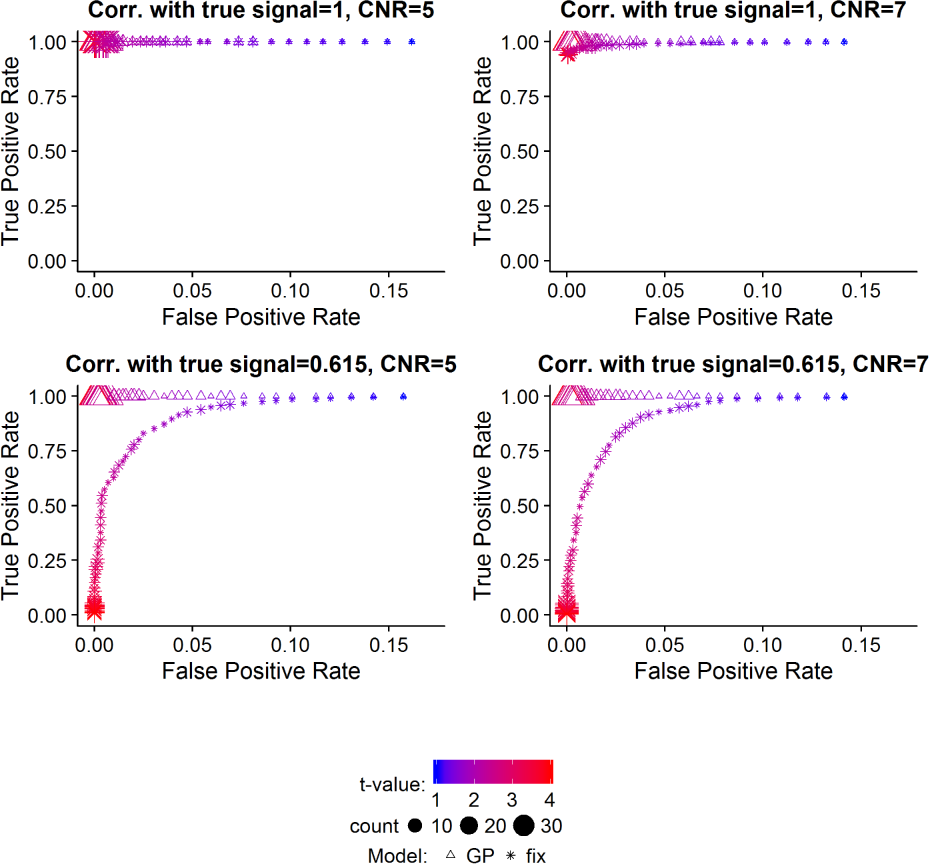
ROC curves for the simulation study. Lengthscale hyperparameter for GP: *l* = 2. Each value of is the average of over 32 simulations. Thresholds for *t*-values: 60 equidistant values between 1 and 4.

**Figure 3:**
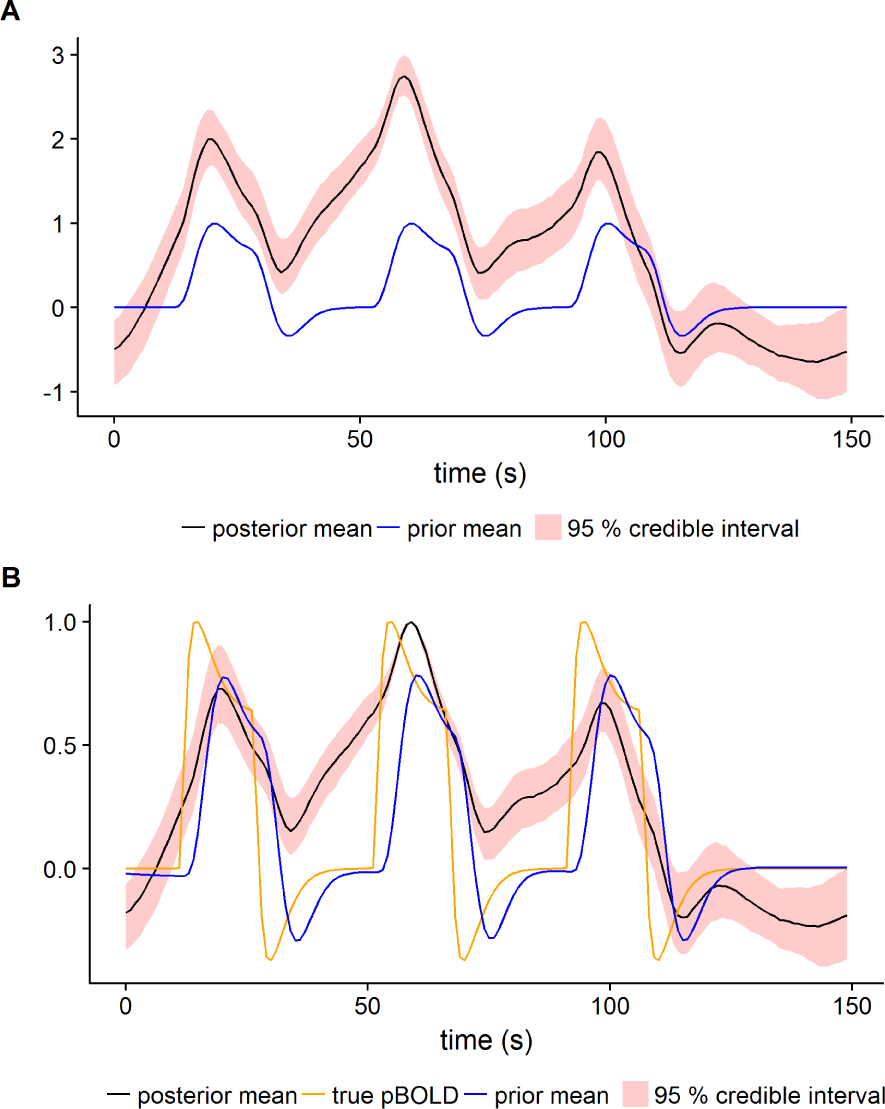
Posterior for predicted BOLD for a simulated parcel. CNR=5, correlation with true signal is 0.615. Lengthscale hyperparameter for GP: *l* = 4. (A) is the Gaussian process **F** in 3 and (B) is the transformed Gaussian process *H*(**F**), see Equation (1) and (2).

**Figure 4:**
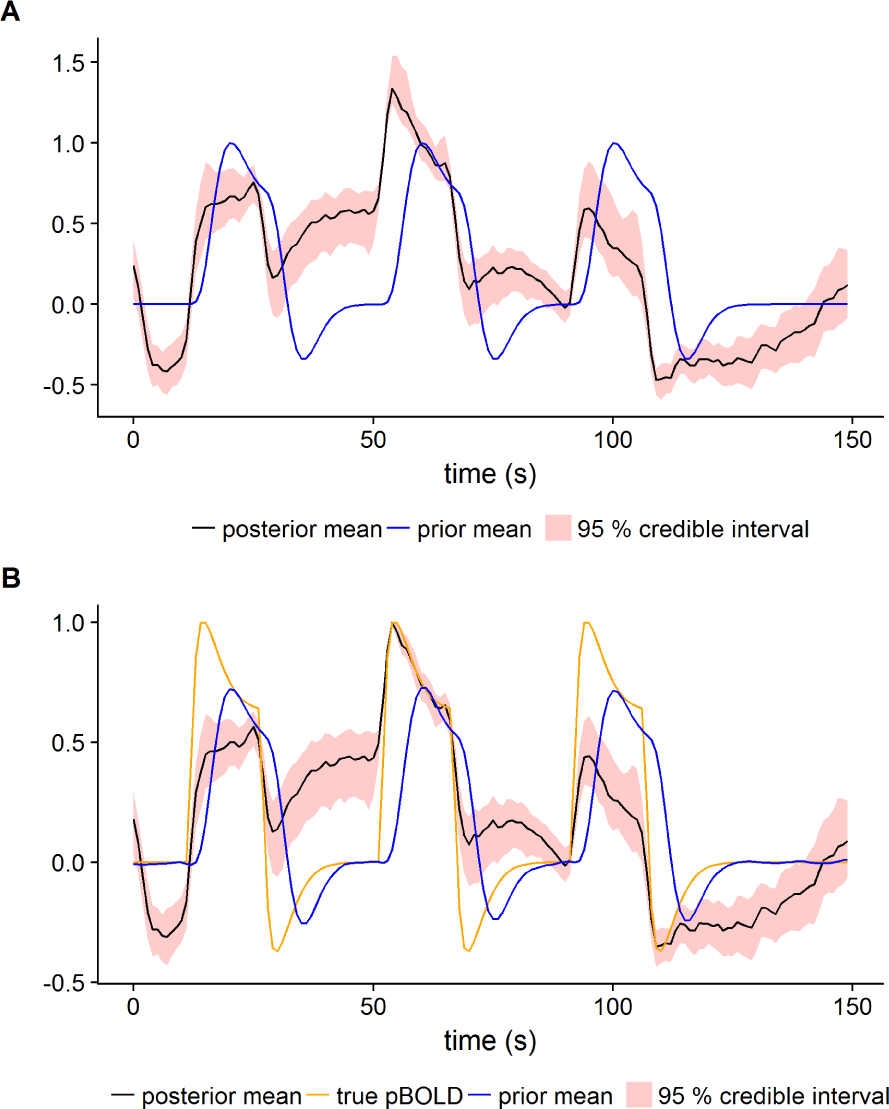
Posterior for predicted BOLD for a simulated parcel. CNR=5, correlation with true signal is 0.615. Lengthscale hyperparameter for GP: *l* = 2. (A) is the Gaussian process **F** in 3 and (B) is the transformed Gaussian process *H*(**F**), see Equation (1) and (2).

We here focus on the word repetition task. The task is to repeat a given word (overt word repetition), in 6 blocks with 30 seconds of activation and 30 seconds of rest. We can thereby expect activation of the language areas of the brain, parts of the motor cortex that correspond to the mouth and tongue (speech production) and the auditory cortex (listening). Our presented results are for two randomly selected subjects: 18716 and 19628.

### 4.2. Preprocessing

The fMRI data were preprocessed using motion correction and 6 mm smoothing. The brain parcellation was performed by registering the ADHD 200 parcel atlas^1^ (Craddock et al., 2012; Bellec et al., 2017) to EPI space, by combining linear T1-MNI and EPI-T1 transformations using FSL. A brain mask is applied in order to remove voxels outside the brain.

In order to be able to compare effect sizes between voxels, the fMRI data is scaled. We use the scaling

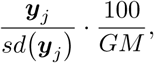

where *y_j_* is observed data for voxel *j*, *sd*(.) is the standard deviation function and *GM* is the global mean of *y_j_*/*sd*(*y_j_*) over all voxels. This scaling ensures that overall the voxels have an average value close to 100 and the same standard deviation. This makes it possible to use effect sizes in terms of percent of the global mean signal, as done in Penny et al. (2005); Sidén et al. (2017). There are 186 parcels and a total of 19,836 voxels for subject 18716. Corresponding numbers for subject 19628 are 179 and 19,304, respectively. Some example parcels and the distribution of parcel size for subject 19628 are shown in Figure 5.

### 4.3. Results

Independent models were fitted to each parcel. An autoregressive model of order 3 was used for the noise process. The same priors were used for all parcels. The same prior hyperparameters were used as in the simulation study except for **F**, which used kernel hyperparameters *l* = 4 and *ω* = 0.1. The prior mean function for **F** was specified to the default HRF in SPM convolved with the paradigm, and was scaled to have zero mean and unit variance.

**Figure 5:**
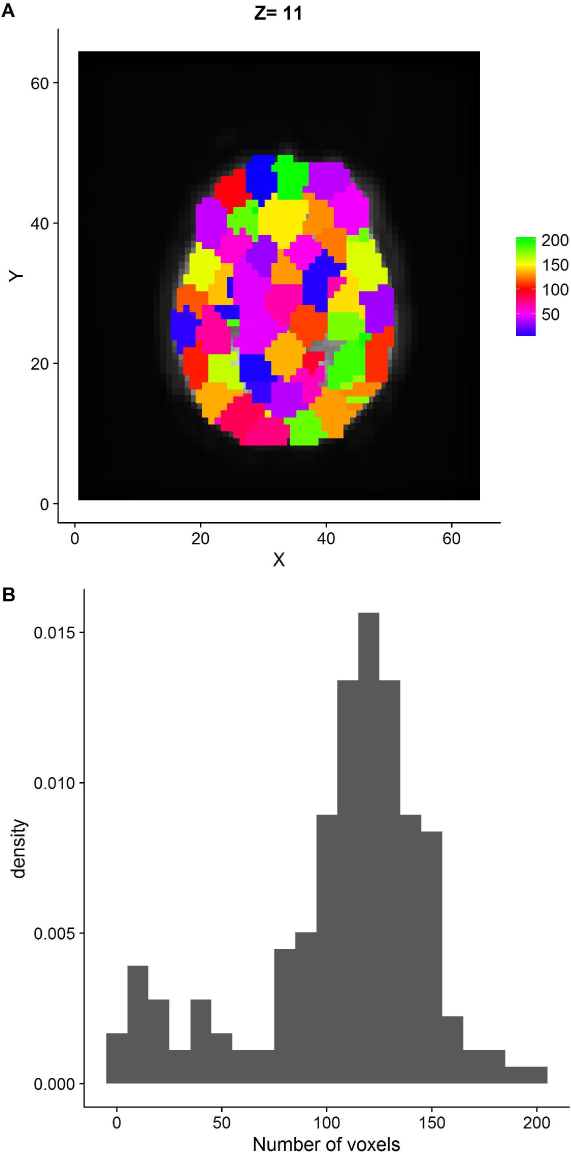
Descriptive statistics for subject 19628. (A) shows all parcels for Z-slice 11. The color specifies parcel belonging. (B) shows the number of voxels in all parcels. Note that our model falls back on the prior mean if there are few active voxels in a parcel, meaning that the model will not break down for small parcels.

Constant, linear, quadratic and cubic trends were included as nuisance variables. Starting values were obtained by using the prior mean for **F**, and all other parameters were initialized using an Cochrane-Orcutt estimation procedure in each parcel. Group-wise ridge regression, using the R-package glmnet (Friedman et al., 2010), was used to obtain the nuisance and the activation parameters. The autoregressive parameters were estimated for the whole parcel using regularized heteroscedastic regression similar to (6). The two steps were iterated until the difference in mean squared error was less then 0.01.

The proposed GP model is compared with three baseline models. In the first model the predicted BOLD is fixated to the prior mean. The second model also uses the temporal derivative of the prior mean, i.e. two basis functions (which is the most common way to allow for a small time shift of the paradigm). The third model uses a smooth FIR approach (Goutte et al., 2000; Ciuciu et al., 2003; Marrelec et al., 2003a; Makni et al., 2008; Vincent et al., 2010) to model the predicted BOLD (see Appendix C for details). The same priors are used for all other parameters.

The posterior was sampled 9000 times, and 3000 samples were discarded as burn in. The remaining samples were thinned out by a factor 6, leaving a final sample of 1000 posterior draws for inference. The parameters **B**, **Γ**, *σ*^2^ and *ρ* showed a good mixing with low autocorrelation. In some parcels, the elements in **F** had high autocorrelation, but the first and second half of the draws gave similar posteriors.

There can be sizeable differences in activation from using a GP prior on the predicted BOLD, compared to using a fixed predicted BOLD. For example, focusing first on subject 19628, Figures 6 and 7 show that the model that estimates the predicted BOLD finds more activity compared to the two first baseline models. For example, the flexible model detects more brain activity in Broca:s language area, which for this subject is close to the brain tumor. According to Figure 8, the flexible GP model also detects stronger brain activity compared to the smooth FIR filter approach.

Figures 9 and 10 show the estimated posterior for the predicted BOLD in the two parcels with most positive activation for subject 19628. Parcel 159 is the yellow cluster in Z-slice 10 and 11 in Figure 6 and has 47 active voxels in total. Parcel 32 is the purple cluster in Z-slice 10 and 11 in Figure 6 and has 47 active voxels in total. Note that the scale of the transformed GP is relative and bounded between -1 and 1, due to the infinity norm. The posteriors have a non-linear behavior, where the amplitude of the peaks and the undershoots vary over time. This form of the posteriors indicate that the data contain important information about the shape of the predicted BOLD, which is not contained in the prior.

Turning now to subject 18716, none of the two first baseline models (using standard basis functions) detected any activity for the given effect size, but our model that estimates the predicted BOLD detected several active voxels, which can be seen in Figure 11. For example, the flexible model detects brain activity in auditory cortex and in motor cortex, not detected by the fix model. As can be seen in Figure 12, the smooth FIR filter approach also detects activity in the motor cortex, but not in the auditory cortex. It should be noted, however, that this activity difference is not caused by a different HR due to the tumor, as the detected activity is on the opposite side of the tumor. The results for the baseline model with two basis functions are not shown.

Figure 13 shows the estimated posterior for the predicted BOLD in one of the parcels with most positive activation for subject 18716. Parcel 32 is the blue cluster in Z-slice 12 in Figure 11 and has 25 active voxels in total. Similar to the posterior predicted BOLD shown for subject 19626, the amplitudes of the peaks and the undershoots are non-stationary.

In order to investigate the effect of the lengthscale hyperparameter, GP models with different lengthscales were estimated. The result is presented in Figure 14. With the shorter lengthscales, the predicted BOLD gets more flexible, and thus finds more activity. Similarly, longer lengthscales restrict the model and the results are more similar to the reference model.

In parcels that lack activity, the posterior predicted BOLD is similar to the prior distribution (not shown).

## 5. Discussion

### 5.1. Model the predicted BOLD instead of the HRF

We have proposed and implemented a new way to model the predicted BOLD for task fMRI. The difference from other models is the direct modeling of the predicted BOLD time series, instead of the HRF, combined with a straight forward measure of activity. The simulation study shows that the model has a good ability to discriminate between active and non-active voxels. The proposed model shows robustness to misspecification in the prior mean function for the predicted BOLD. This is a desirable feature, since it is likely that a model with a fix predicted BOLD will not be correct for the whole brain or across subjects. Our proposed model gives the researcher a framework to approach problems related to the HR. For example, for group studies (where all data are transformed to a standard space), the predicted BOLD in one parcel can be compared across subjects. Also, the existence of explicit activity parameters makes it easy to construct PPMs or *t*-maps.

The non-linear aspect of the hemodynamics can be captured with the GP model. It is interesting to study the properties of the posterior for the predicted BOLD, see Figures 9, 10 and 13. Compared to the prior mean function, the major difference for the posterior is the timevarying amplitude of the peaks and the undershoots. This feature seems to be crucial to find the additional activity compared to the two baseline models and the smooth FIR approach. The described feature is not easily incorporated into traditional GLM approaches. Parametric modulation of the HRF can be used in the LTI context, to obtain hemodynamic features that are non-stationary, but this approach comes with two problems. First, a proper modulator must be chosen. Second, the non-stationarity is assumed to be be known and fix across the brain given the modulator. Our approach handles the non-stationarity in an unsupervised manner, and can of course use a prior mean function that depends on a problem specific modulator.

### 5.2. Computation

The model is implemented in the R programming language, and the Markov Chain Monte Carlo (MCMC) code is not optimized for speed, except for the CPU parallelization of models across parcels. The computations for our model are therefore rather time consuming, but there are several options to reduce the computational time. One option is to use other inference methods, such as VB inference. Another option is to use GPUs (graphics cards) for parallelization, particularly if there are a relatively large number of parcels.

**Figure 6:**
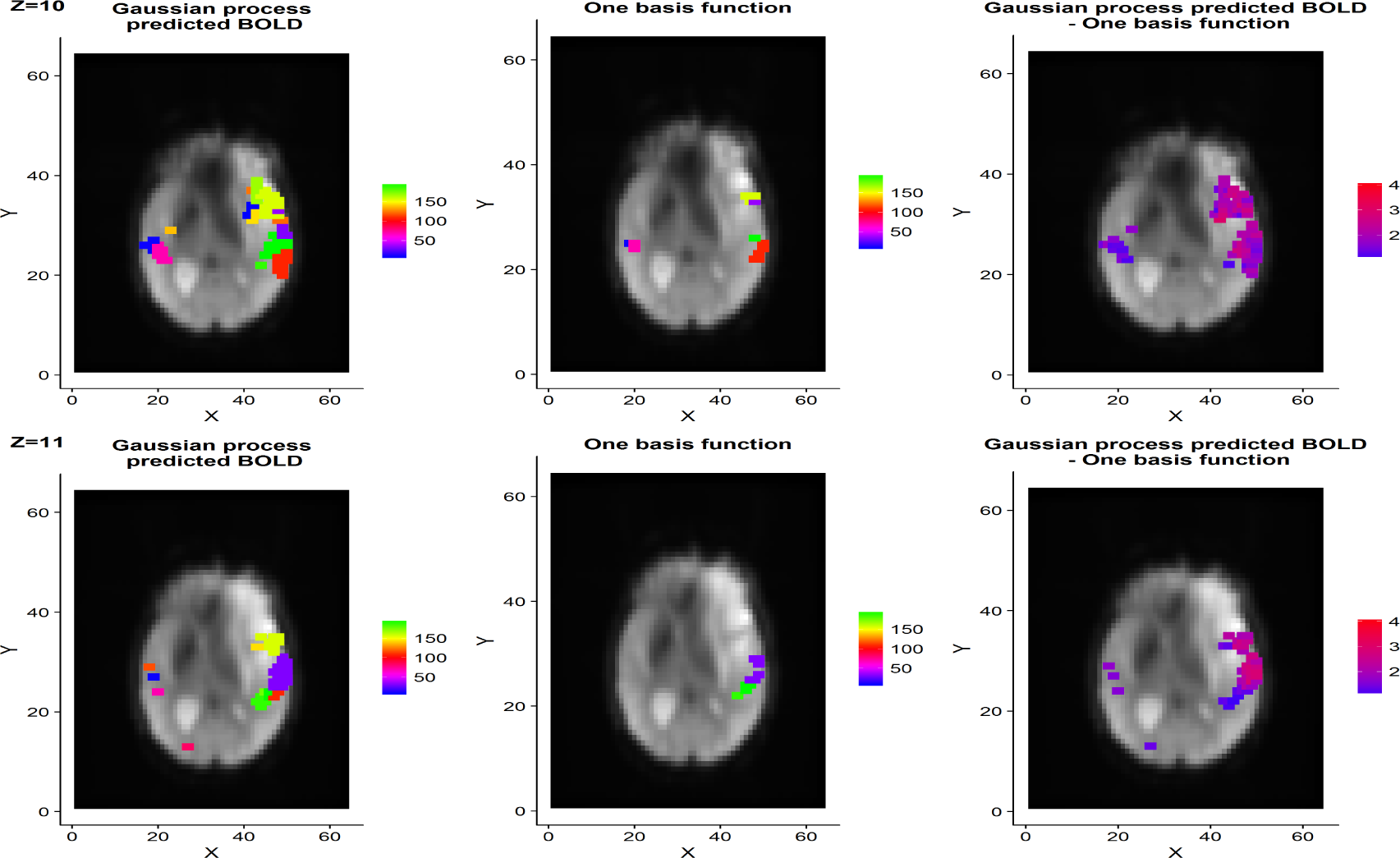
Example slices with Bayesian *t*-ratios for subject 19836. The activity maps are thresholded at *t* ≥ 4 for a test that tests the effect size 0.25. The color specifies parcel belonging for active voxels. The rightmost column shows the differences in t-ratios, thresholded such that only values fulfilling |*t*_1_ – *t*_2_| > 1 are shown. Our flexible model clearly detects more activity, compared to the fix predicted BOLD model. Top row: the flexible model detects more brain activity in Broca1s language area, which for this subject is close to the brain tumor. Bottom row: the flexible model finds brain activity in the visual cortex and bilateral activation of the auditory cortex, which the fix model struggles to detect.

**Figure 7:**
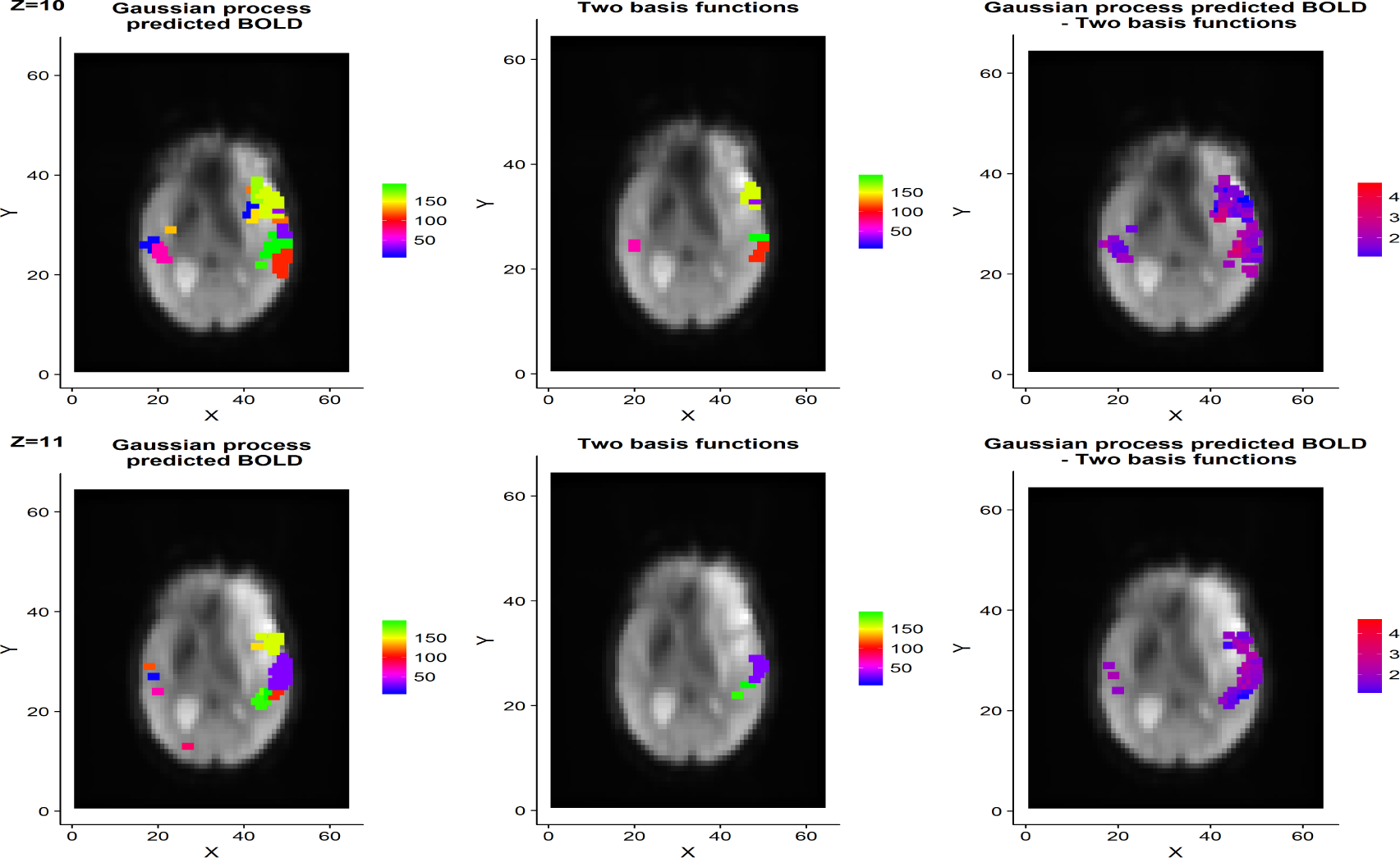
Example slices with Bayesian *t*-ratios for subject 19836. The activity maps are thresholded at *t* ≥ 4 for a test that tests the effect size 0.25. t-ratios for the baseline model is created with *b* for the first basis. The color specifies parcel belonging for active voxels. The rightmost column shows the differences in t-ratios, thresholded such that only values fulfilling |*t*_1_ – *t*_2_| > 1 are shown. The differences between the flexible model and the fix model are very similar to Figure 6, where a single basis function was used.

**Figure 8:**
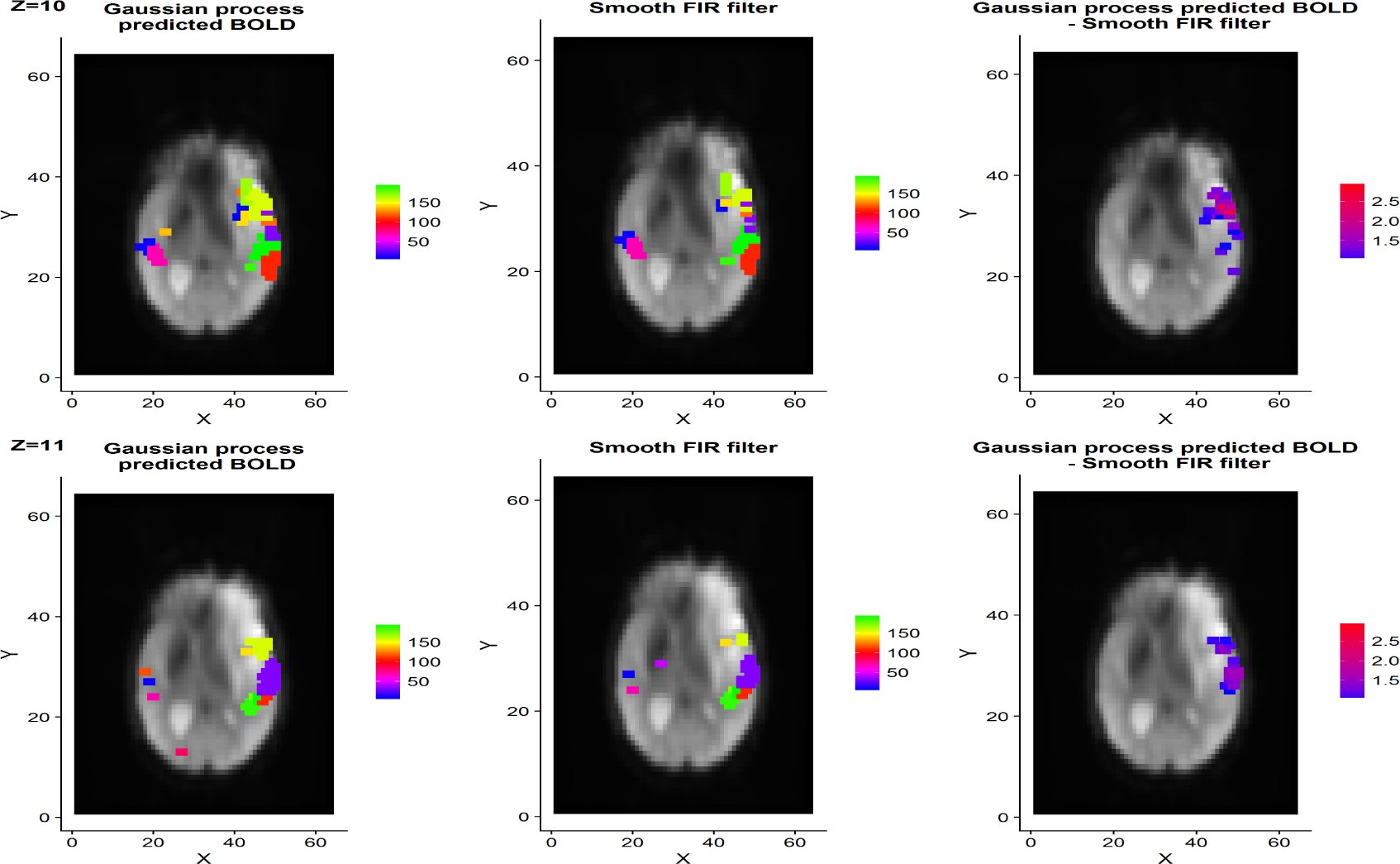
Example slices with Bayesian *t*-ratios for subject 19836. The activity maps are thresholded at *t* ≥ 4 for a test that tests the effect size 0.25. The color specifies parcel belonging for active voxels. The rightmost column shows the differences in t-ratios, thresholded such that only values fulfilling |*t*_1_ − *t*_2_| > 1 are shown.

**Figure 9:**
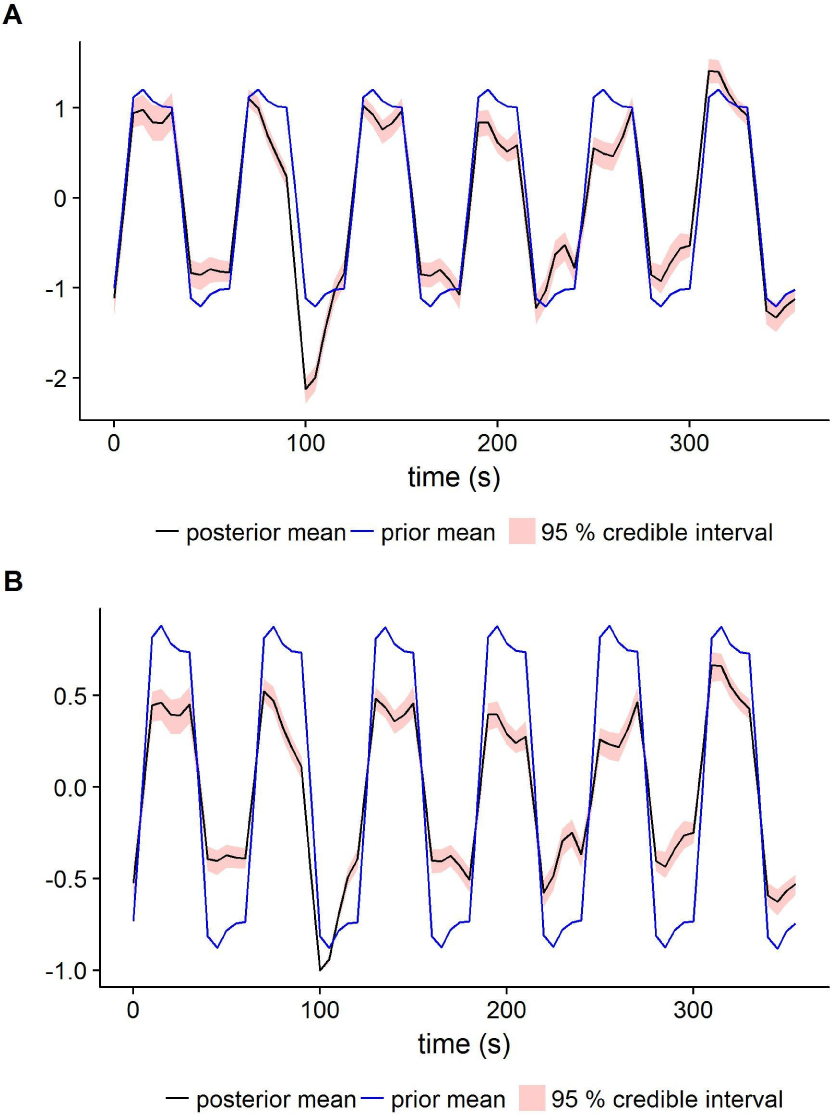
Estimated predicted BOLD for subject 19836 and parcel 159. (A) is the Gaussian process **F** in (3) and (B) is the transformed Gaussian process *H*(**F**), see Equation (1) and (2).

**Figure 10:**
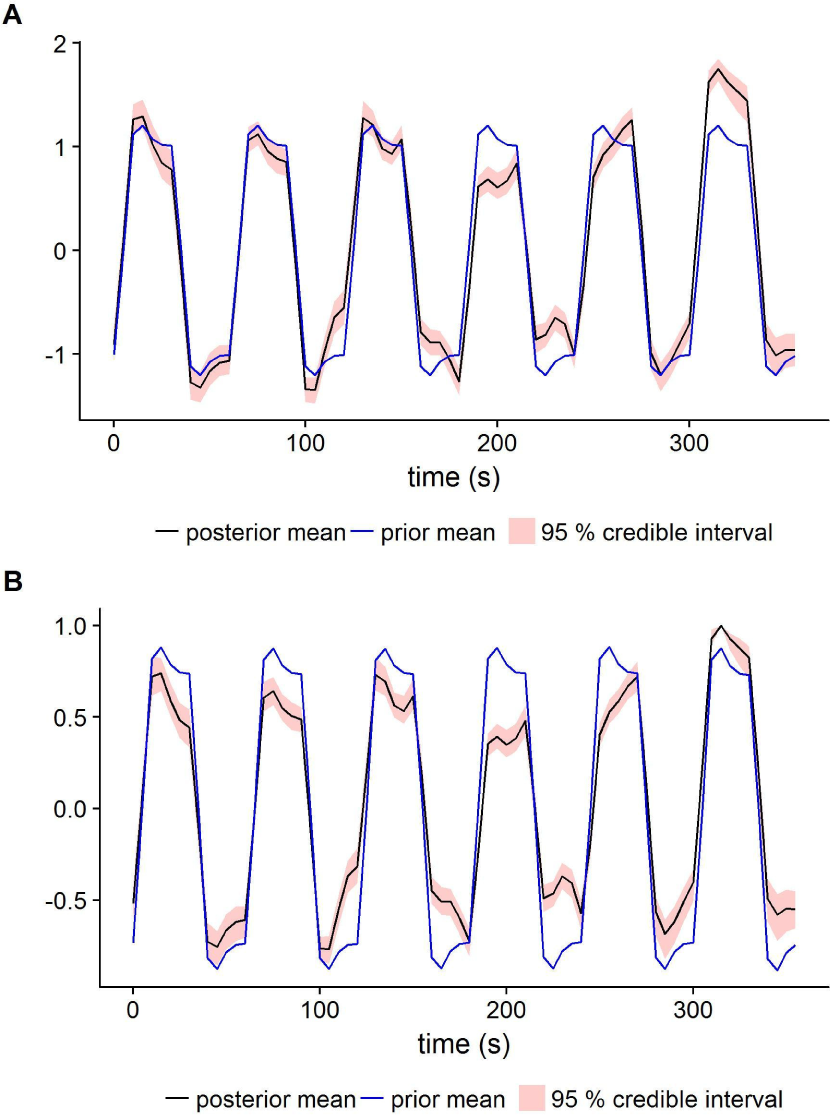
Estimated predicted BOLD for subject 19836 and parcel 32. (A) is the Gaussian process **F** in (3) and (B) is the transformed Gaussian process *H*(**F**), see Equation (1) and (2).

**Figure 11:**
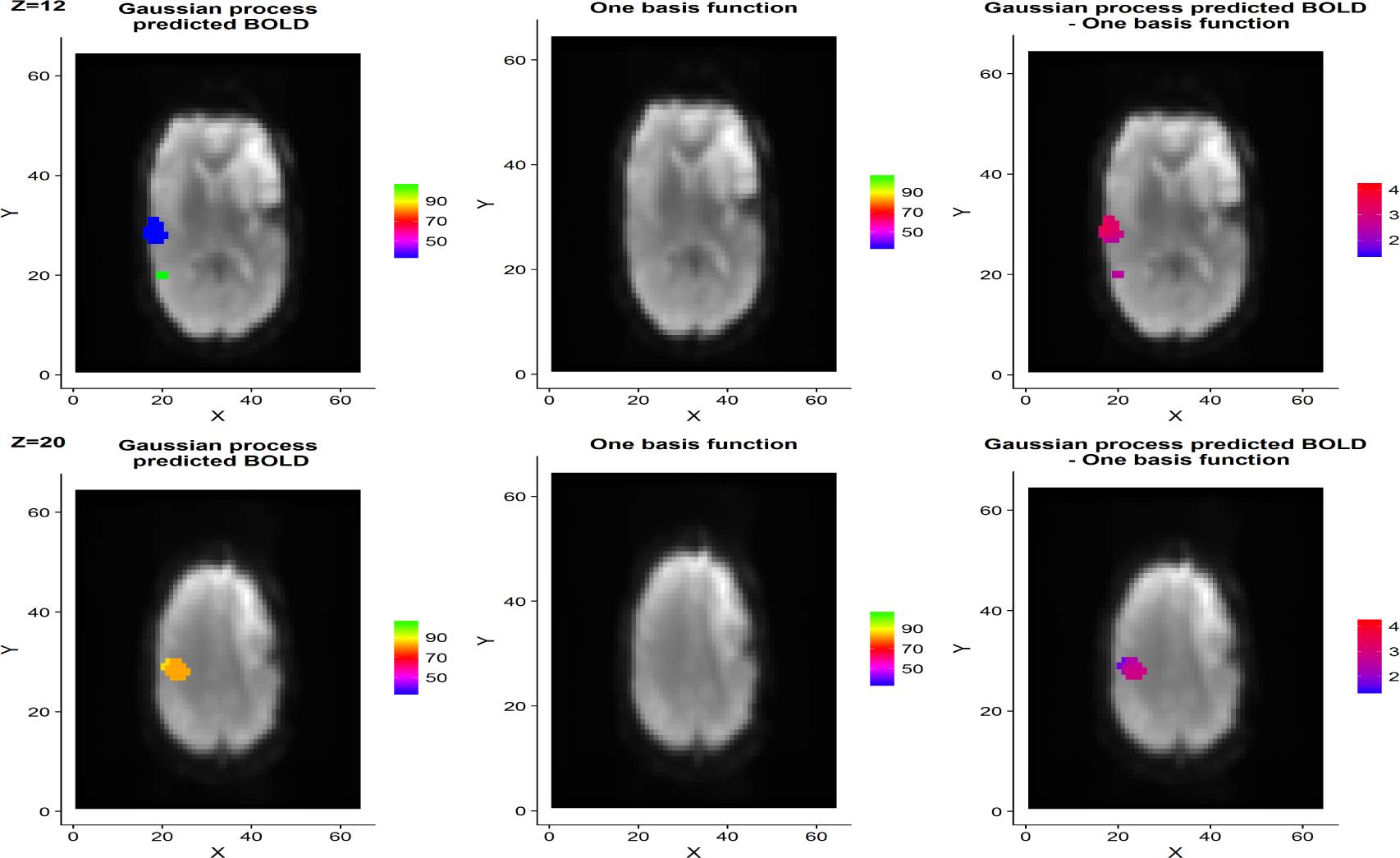
Example slices with Bayesian *t*-ratios for subject 18716. The activity maps are thresholded at *t* ≥ 4 for a test that tests the effect size 0.25. The color specifies parcel belonging for active voxels. The rightmost column shows the differences in t-ratios, thresholded such that only values fulfilling |*t*_1_ – *t*_2_| > 1 are shown. Top row: the flexible model finds brain activity in the auditory cortex, which is not found using the fix model. Bottom row: the flexible model finds brain activity in the motor cortex (generated by speech production), which is not found using the fix model.

**Figure 12:**
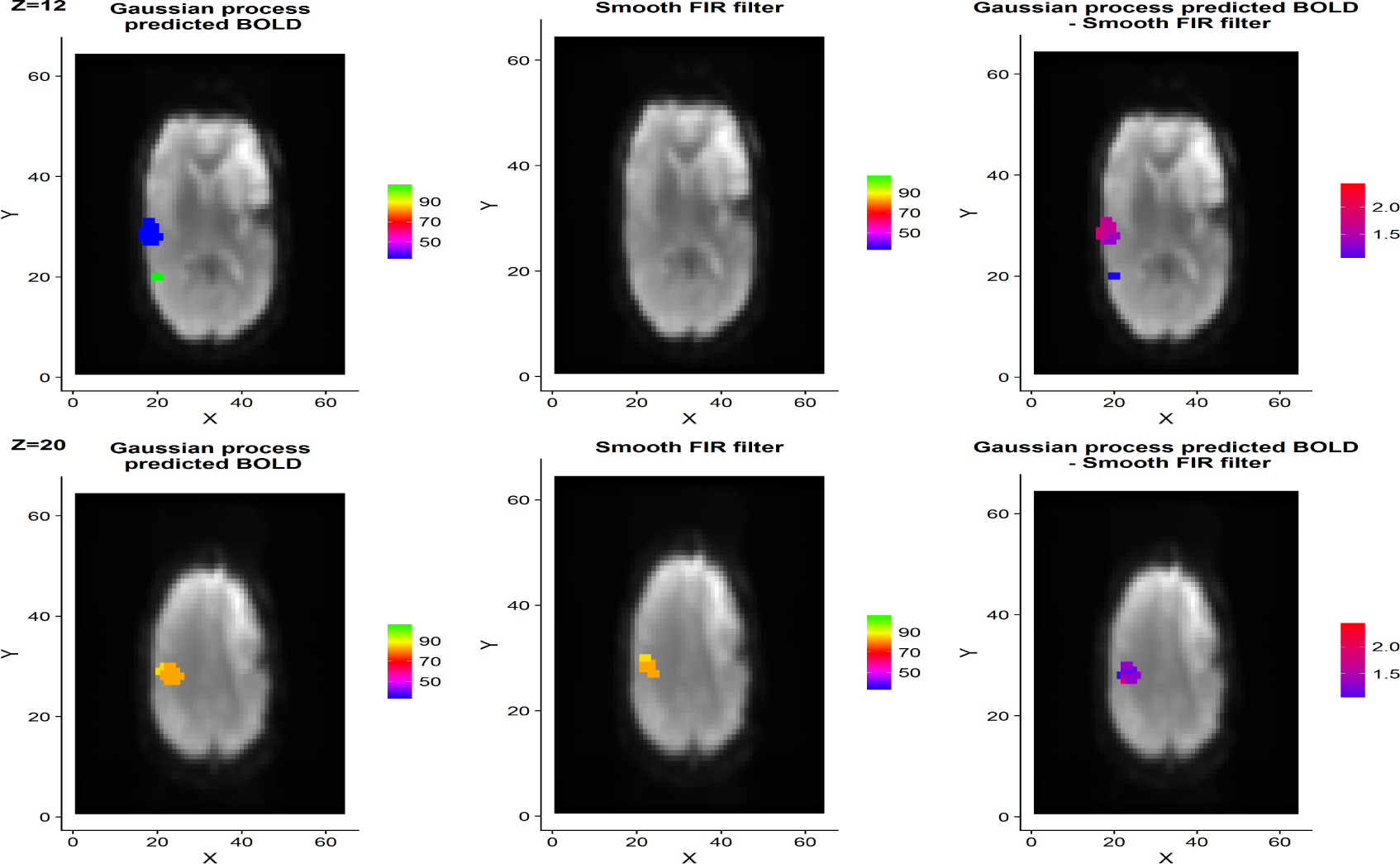
Example slices with Bayesian *t*-ratios for subject 18716. The activity maps are thresholded at *t* ≥ 4 for a test that tests the effect size 0.25. The color specifies parcel belonging for active voxels. The rightmost column shows the differences in t-ratios, thresholded such that only values fulfilling |*t*_1_ − *t*_2_| > 1 are shown.

**Figure 13:**
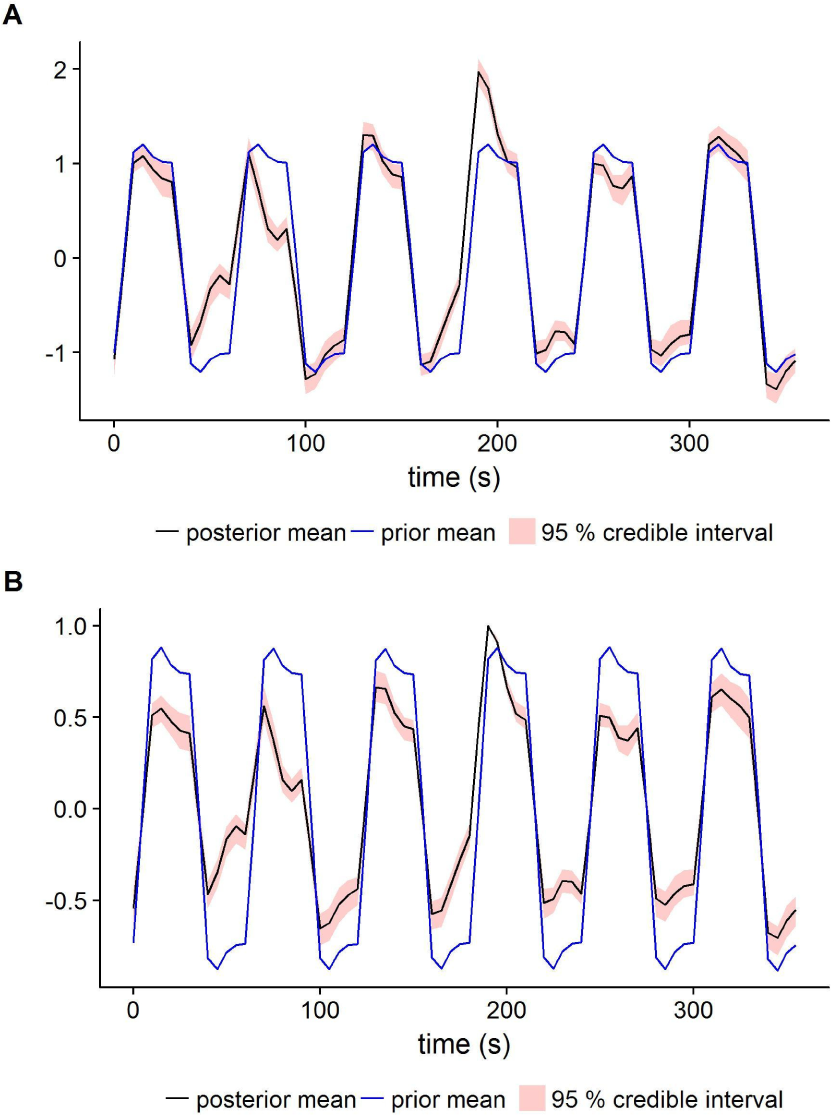
Estimated predicted BOLD for subject 18716 and parcel 32. (A) is the Gaussian process **F** in (3) and (B) is the transformed Gaussian process *H*(**F**), see Equation (1) and (2).

### 5.3. Multiple comparisons

In contrast to frequentist methods, there is no consensus in the fMRI field regarding if and how to correct for multiple comparisons for PPMs. In frequentist hypothesis testing the null hypothesis is normally that the parameter representing the brain activity is 0, but using an effect size threshold of 0 for PPMs often leads to activation in a very large portion of the voxels (even for strict probability thresholds for the PPMs). In this paper we have mainly focused on differences between fix and flexible predicted BOLD models, using voxel inference and an effect size threshold of 0.25. One ad-hoc approach to correct for multiple comparisons is to calculate a Bayesian *t*or *z*score for each voxel, and then apply existing frequentistic approaches for multiple comparison correction (e.g. Gaussian random field theory). This approach is for example used in the FSL software.

### 5.4. Applications

There are several possible applications of our proposed model. As demonstrated in this paper, a potential application is in clinical fMRI, where fMRI can be used to map out important brain areas prior to tumor surgery. The HR may be different close to a tumor, and our flexible model can then be used to detect more brain activity, and thereby potentially lead to a better treatment plan. Other cases where the HR may be different include young subjects (Richter and Richter, 2003), subjects with epilepsy (Jacobs et al., 2008) and subjects with stroke (Bonakdarpour et al., 2007). Our model can also be used to automatically handle cases where a subject fails to perform one or several events, or where a subject occasionally struggles with the timing of the experiment (adding a temporal derivative can only account for a global shift in time, and not local time shifts). As mentioned in the introduction, our model can also pick up variations in the strength of the BOLD response, while virtually all other models see the stimulus as a fixed effect (Westfall et al., 2016).

### 5.5. Future work

A natural extension of the model is to do inference for the GP hyperparameters. Since the model uses a nonGaussian likelihood this is, however, non-trivial. In the MCMC case the methods presented in Filippone and Girolami (2014); Murray and Graham (2016) could be used, where pseudo marginal inference is employed together with an unbiased estimate of the intractable marginal likelihood for the GP.

The prior mean is a quite severely misspecified in the simulation in Section 3, and the GP model cannot fully capture the true underlying signal, see Figures 3 and 4, even though it captures sufficient variation to be able to distinguish between active and non-active voxels. Other kernels could perform even better in this scenario. For block paradigms, locally periodic kernels are a natural candidate, see for example Duvenaud (2014) for a discussion. Those kernels could introduce a positive periodic correlation that decays with the distance; paradigm blocks near in time will be more correlated compared to blocks further away in time. Another alternative is use the prior mean function *m*(*t*) and the derivative *∂m*(*t*)/*∂t* as covariates in a smooth kernel. This would give a behavior similar to locally periodic kernel, but can also handle event related data. The number of hyperparameters grows with more complex kernels, and it is hard to manually specify these parameters. To efficiently use more complex kernels, inference for the hyperparameters is important, either using an estimate of the posterior mode or sampling via MCMC. Even more sophisticated kernels could be used, such as spectral mixture kernels (Wilson and Adams, 2013) or deep kernels (Wilson et al., 2016). Such kernels are very expressive and can find complicated patterns given sufficient data. However, the proposed inference methods are adapted for the classical GP regression model *y = f* (*x*)+*ε*, where *f* is a GP, and not as a part of a multivariate time series regression model. The problem boils down to finding data driven features that can be used in some kernel, in order to explain the prior correlation in **f** in a good way for a given parcel.

Another possible future direction is to use more sophisticated priors on **B** and **Γ**. Here variable selection would be appropriate, as it could reduce overfitting and improve the model fit. Interesting selection methods could be spike and slab priors or horseshoe priors. Due to the spatial nature of the data, spatial priors would also be an interesting choice. For example, a GP prior could be placed over **B** to impose a spatial smoothness, e.g. using a Matern kernel. This would result in models similar to those in (Luttinen and Ilin, 2009) and (Wilson et al., 2012). Sidén et al. (2017) use sparse precision matrices to model spatial dependencies in whole brain task fMRI data, and derive both fast MCMC and VB methods. Those ideas could be incorporated into the proposed model.

**Figure 14:**
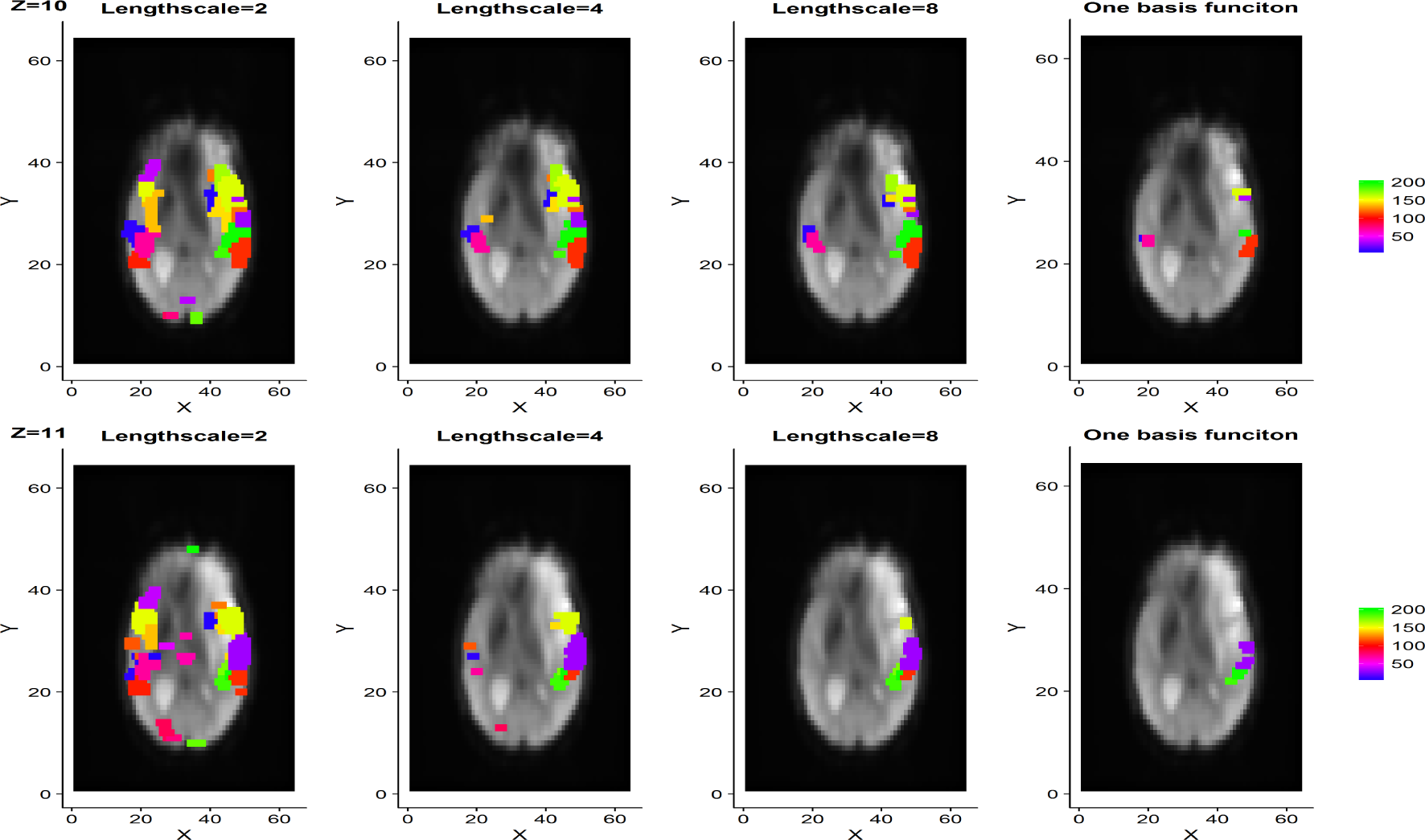
Comparison of different lengthscales for the GP model. Example slices with Bayesian *t*-ratios for subject 19836. The activity maps are thresholded at *t* ≥ 4 for a test that tests the effect size 0.25. The color specifies parcel belonging for active voxels. Clearly, decreasing the lengthscale leads to a more flexible predicted BOLD, which may lead to overfitting.

The suggested model is made for single subject data, and in many cases joint inference for many subjects is desirable. A hierarchical model could be used here, with random effects for the **B** and **F** parameters. Assume that all subjects have been transformed to the same space with the same parcellation. The prior for **f***_m_* for a given parcel could then be expressed as

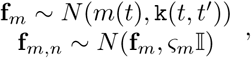

where *n* is subject index and *ς_m_* is the random effects variance. **B** can be modeled in the same way. This construction is similar to the within subject models used in (Chaari et al., 2012, 2016; Albughdadi et al., 2016), but a difference is that they have a random effect on the HRF filter and the hierarchy is over parcels and voxels.

## 6. Conclusion

We have proposed a novel framework for modeling the hemodynamics in task fMRI. The new model is shown to more accurately detect brain activity compared to traditional parametric and nonparametric LTI models. We model the predicted BOLD directly with a GP prior, as a part of larger time series regression model. We also introduce an identifying transformation that solves the challenging identification problem present in bilinear models in the JDE context. This can be done due to problem specific constraints related to the hemodynamics. Our new framework gives researchers the opportunity to ask new kinds of questions related to hemodynamics, especially with regard to non-linear effects.

## Acknowledgments

This work was funded by Swedish Research Council (Vetenskapsradet) grant no. 2013-5229.

## Appendix A: Distributions

The density function for a matrix normal distribution

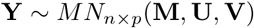

is of the form

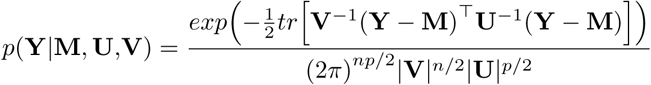

Parameters:

- **M**: location, real *n* × *p* matrix
- **U**: scale, positive-definite real *n × n* matrix (dependencies over observations)
- **V**: scale, positive-definite real *p × p* matrix (dependencies over variables)

The matrix normal distribution is related to the multivariate normal distribution in the following way:

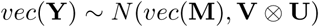

## Appendix B: Gibbs sampling

### Sampling *ρ*

The autoregressive parameters are sampled using the following formulation

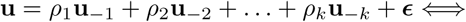

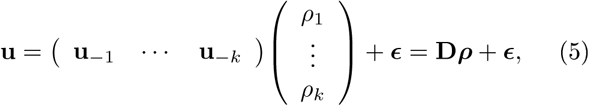

where *u_*_* = *vec*(**Y***_*_* − **F***_*_***B** − **Z***_*_***Γ**) is used to calculate **u**, given that **B**, **F***_*_* and **Γ** are known, **D** is a matrix of size *JT* × *k*. **u** and **D** must be updated in every iteration since **B**, **F** and **Γ** also are updated in every iteration. Let **Σ** = **Ω** ⊗ П*_T_*, then *ε* ~ *N* (**0**, **Σ**). It is clear from (5) that the full conditional posterior for *ρ* can be obtained using standard formulas for univariate regression with heteroscedastic variance, and the full conditional posterior is given by

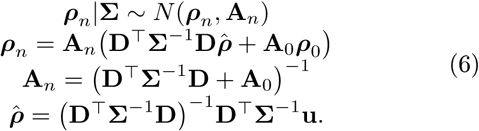

To make sure that *ρ* is in the stationary region, draws from *ρn*|**Σ** ~ *M N* (*ρn*, **A***_n_*) are discarded until a draw that is inside the stationary region is obtained.

### Sampling B, Γ and *σ*^2^

In order to simplify sampling of **B** and **Γ** in model (3) the following formulation is used

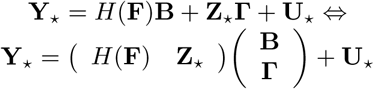

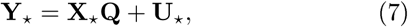

where **X***_*_* is of the size *T_*_* × (*M + P*) and **Q** is of the size (*M + P*) × *J*. In order to use the standard multivariate regression formulas in (7), pre-whitening is used. Let Φ*_C_* (*L*) be the column-wise lag polynomial from time series analysis, i.e. Φ*_C_* (*L*) = 1 − *ρ*_1_*L* − *ρ*_2_*L*^2^ − *…* − *ρ_k_L^k^*. The first *K* constant observations in **Y**_0_, **F**_0_ and **Z**_0_ are used for obtaining the first *K* observations of 
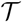
. This results in a new regression formulation

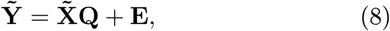

where **Y˜** = Φ*_C_* (*L*)**Y**, **X˜** = Φ*_C_* (*L*)**X** and **E** = **U˜** = Φ*_C_* (*L*)**U**.

Given the parcel constant parameters **F**, *ρ*, and the regression parameters **Q**, the likelihood *p* **Y˜**, **X˜**|*σ*^2^, **Q** is independent over voxels, which implies that inference for each element in *σ*^2^ is performed by regressions of the form: **y˜***_j_* = **X˜q***_j_* + *ε_j_*, where **y˜***_j_* is the j:th column of **Y**˜ and **q***_j_* is the j:th column of **Q**. The full conditional posterior for 
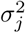
 is an Inverse-gamma distribution, which is easily obtained from standard formulas for univariate regression

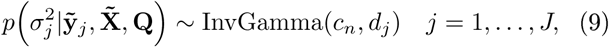

where

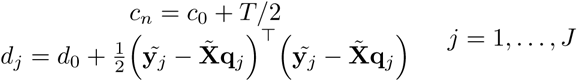

Using standard formulas for multivariate regression with conjugate priors, the full conditional posterior for *vec*(**Q**) is a multivariate normal distribution. The likelihood function for (8) is described by Press (1982). Let 
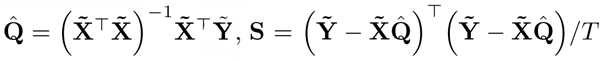
 and *vec*(**Q**)=*q*. Using standard manipulations, the likelihood can be written as

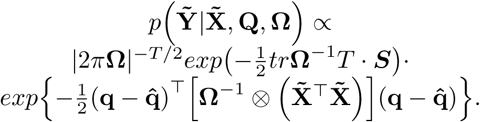

The prior for **Q** is now expressed as

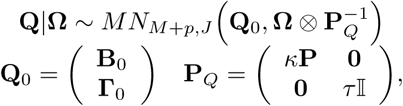

where **Q**_0_ has the same size as **Q** and **P***_Q_* has size (*M + P*)× (*M + P*). Using standard formulas for multivariate regression, the full conditional posterior for *Q* is then given by

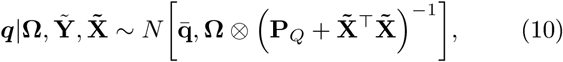

where

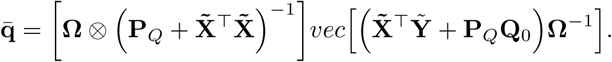

### Sampling F

To simplify the sampling of **F**, we reformulate the model as

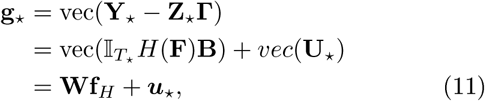

where **W** = (**B***^T^* ⊗ П*_T_\** is of size *JT_*_* × *T_*_M* and **f***_H_* = vec(*H*(**F**)) is of size (*JT_*_*) × 1. Now, Equation (11) is transformed with a lag polynomial Φ*_R_*(*L*) in a row-wise manner. Φ*_R_*(*L*) has the same functional form as Φ*_C_* (*L*), but operates independently on each voxel time series. The first *T_*_* rows will first be transformed, followed by transformation of the next *T_*_* rows, until all rows have been transformed. The transformation results in

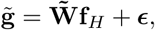

where **W**˜ is of size *JT* × *T_*_M*, **g**˜ and *ε* are both of size (*JT*) × 1. The likelihood for **f***_H_* is given by

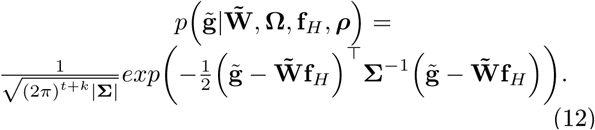

Note that **F** enters the Gaussian likelihood in a non-linear way, which means that the full conditional posterior is not available in closed form. We use elliptical slice sampling Murray et al. (2010) to sample from the posterior of **F**. Elliptical slice sampling is a slice sampling technique which is particularly suitable for Gaussian process models with non-Gaussian likelihoods.

## Appendix C: Smooth FIR model for predicted BOLD

The proposed model is also compared with a smooth FIR model. In order to make the comparison as fair as possible, the FIR model is formulated as

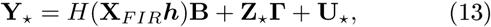

where **X***_FIR_* is the standard FIR design matrix (where stimuli are organized column wise) with *K × M* columns. *K* is the filter length, and *h* is the filters for all stimuli stacked in one vector of size *KM* × 1. *h* has independent and identical GP priors for each stimuli. Since *h* contains much fewer parameters than **F**, the scale factor of the kernel is specified to a higher value compared to **F**, in order to increase flexibility. The filters in *h* is estimated using elliptical slice sampling, in the same way as **F** is estimated in our model. The inference for the other parameters is not changed. The prior mean function for *h* was specified to the common double gamma HRF, and the√kernel hyper-parameters were specified to *l* = 3 and 
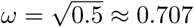
. The endpoints of the filter were constrained to the corresponding prior mean values.

## Footnotes

1 https://www.nitrc.org/frs/?group_id=427

